# MicroRNA-223 protects neurons from degeneration in Experimental Autoimmune Encephalomyelitis

**DOI:** 10.1101/430777

**Authors:** Barbara Morquette, Camille A. Juźwik, Sienna S. Drake, Marc Charabati, Yang Zhang, Marc-André Lécuyer, Dylan Galloway, Aline Dumas, Omar de Faria, Mardja Bueno, Isabel Rambaldi, Craig Moore, Amit Bar-Or, Luc Vallières, Alexandre Prat, Alyson E. Fournier

## Abstract

Multiple sclerosis (MS) is an autoimmune disease characterized by demyelination and neurodegeneration in the brain, spinal cord and optic nerve. Neuronal degeneration and death underlie progressive forms of MS and cognitive dysfunction. Neuronal damage is triggered by numerous harmful factors in the brain that engage diverse signalling cascades in neurons thus therapeutic approaches to protect neurons will need to focus on agents that can target broad biological processes. To target the broad spectrum of signaling events that mediate neurodegeneration in MS we have focused on non-coding small microRNAs (miRNAs). microRNAs are epigenetic regulators of protein expression, targeting messenger RNAs (mRNAs) and inhibiting their translation. Dysregulation of miRNAs has been described in many neurodegenerative diseases including MS. In this study we identified two miRNAs, miR-223-3p and miR-27a-3p, that were upregulated in neurons in the experimental autoimmune encephalomyelitis (EAE) mouse model of CNS inflammation and in active MS lesions. Overexpression of miR-27a-3p or miR-223-3p protected dissociated cortical neurons from degeneration in response to peripheral blood mononuclear cell conditioned media (PBMC-CM). Introduction of miR-223-3p *in vivo* in mouse retinal ganglion cells (RGCs) protected RGC axons from degeneration in the EAE model. By *in silico* analysis we found that mRNAs in the glutamate receptor (GluR) pathway are enriched in miR-27a-3p and miR-223-3p targets. Antagonism of the GluR pathway protected neurons from PBMC-CM-dependent degeneration. Our results suggest that miR-223-3p and miR-27a-3p are upregulated in response to inflammation to mediate a compensatory neuroprotective gene expression program that desensitizes neurons to glutamate by downregulating mRNAs involved in GluR signalling.

## Introduction

Multiple Sclerosis (MS) is a complex disease resting at the neuro-immune interface. Peripheral immune cells invade the central nervous system (CNS), leading to inflammation and focal demyelinating lesions (Compston and Coles, 2008). Most cases manifest with a relapsing-remitting (RRMS) clinical disease course, with about half of the cases developing into progressive clinical deterioration. Approximately 15% of MS cases manifest as a progressive disease without remission, termed primary progressive MS (PPMS) (Antel *et al*., 2012). While demyelination correlates strongly with RRMS, axonal degeneration and neuronal death correlate with progressive forms of the disease (Bjartmar *et al*., 2000; Wujek *et al*., 2002). The presence of diffuse axonal damage throughout MS patient tissue implies that degeneration occurs even in the absence of acute inflammation (Nikic *et al*., 2011; Criste *et al*., 2014). Most current MS therapies are immunomodulatory and therapies targeting the neurodegeneration underlying progressive and sustained clinical disability in MS remains an area of unmet clinical need (Kasarello *et al*., 2017).

Multiple factors including cytokines, complement, free radicals, nitric oxide, proteases, excess glutamate and calcium contribute to a harmful environment for vulnerable demyelinated axons in MS (Trapp *et al*., 1998a; Yong *et al*., 2007; Gonsette, 2008). Similarly neuronal damage in MS is triggered by numerous intracellular molecular cascades (Yang *et al*., 2015). Interventions aimed at individual molecular targets may offer only partial neuroprotection and targeting programs of gene expression represents a viable strategy for the development of neuroprotective agents (Yong *et al*., 2007; Gonsette, 2008). microRNAs (miRNAs) are evolutionarily conserved short non-coding RNAs that regulate gene expression in healthy and pathological conditions (Bartel, 2004). Mature miRNAs regulate programs of gene expression by binding to the 3’ untranslated regions (3’UTR) of target messenger RNAs (mRNAs) and inhibiting protein translation or initiating mRNA degradation of multiple mRNA transcripts simultaneously (Grimson *et al*., 2007; Friedman *et al*., 2009; Quinlan *et al*., 2017). A major advantage of targeting miRNAs is therefore that one can modulate a pool of genes rather than a single gene and these pools of genes can be functionally related.

In MS, miRNA profiles are altered in immune cells, cerebrospinal fluid (CSF) and within CNS lesions (Junker *et al*., 2009; Jr Ode *et al*., 2012; Miyazaki *et al*., 2014). miRNAs are also regulated in experimental autoimmune encephalomyelitis (EAE), an animal model of CNS inflammation that exhibits CNS pathology including ascending paralysis and motor neuron damage and death (Bannerman *et al*., 2005; Vogt *et al*., 2009; Hoflich *et al*., 2016). The visual system is also affected in EAE with loss of retinal ganglion cells (RGCs) and an associated optic nerve pathology that includes immune cell infiltration, demyelination and glial activation (Shindler *et al*., 2008; Quinn *et al*., 2011; Horstmann *et al*., 2013).

Using the EAE model miRNAs have been profiled in purified cell types including immune cell subsets, oligodendrocytes and neurons providing valuable information about molecular substrates that may be targeted to manipulate cell specific biologies (Jr Ode *et al*., 2012; Lewkowicz *et al*., 2015; Juzwik *et al*., 2018). We profiled miRNA expression in lumbar motor neurons and retinal neurons in EAE to identify molecular substrates that may be targeted to promote neuroprotection (Juzwik *et al*., 2018). Here, we study two miRNAs, miR-223-3p and miR-27a-3p that are upregulated in motor neurons in mice subjected to EAE. We report that these miRNAs are regulated in active lesions from post mortem MS patient tissue and that overexpression of these miRNA mediate neuronal protection in part through regulation of glutamate receptor signalling.

## Materials and Methods

### Preparation of peripheral blood mononuclear cell conditioned media (PBMC-CM)

Animal procedures were approved by the Montreal Neurological Institute Animal Care and Use Committee, following the Canadian Council on Animal Care (CCAC) guidelines. Blood from adult Sprague-Dawley rats was collected by cardiac puncture. Rat PBMCs were separated by Ficoll-Paque density centrifugation and cultured in Ultraculture serum-free medium (Lonza) containing 1% penicillin-streptomycin for 4 days in vitro (DIV) at 37°C, 5% CO 2 (Pool *et al*., 2011; Pool *et al*., 2012). Control media was Ultraculture serum-free medium (Lonza) containing 1% penicillin-streptomycin incubated without cells for 4 DIV. Individual batches of PBMCCM were titered by determining the dose that mediated 50% neurite outgrowth inhibition; this was defined as the 1X treatment concentrations. Dose curves of conditioned media were then applied at 0.5X – 4X doses by diluting PBMC-CM into volumes of control CM such that all neurons were treated with equal volumes of CM.

### Degeneration assays

Cortical neurons from wildtype (WT) C57Bl/6 mice were prepared as previously described (Juzwik *et al*., 2018; Pare *et al*., 2018). Cortical neurons were seeded in 48-wells at 80 000 cells/well and aged 4 DIV prior to addition of PBMC-CM. Treated neurons were fixed with 4% paraformaldehyde (PFA)/20% sucrose and stained with anti-βIII tubulin antibody (Covance), Hoechst 33342 dye (Sigma-Aldrich) and fluorescent secondary antibodies. Apoptotic cells were identified by TUNEL staining (In Situ Cell Death Detection Kit, Roche). Automated image acquisition and analysis were performed using ImageXpress and the Multi Wavelength Cell Scoring (MWSC) module of MetaXpress (Molecular Devices) to determine the number of neuronal cell bodies by overlapping βIII tubulin and Hoechst staining. Fixed and imaged neurons were analyzed for loss of neurite processes using ImageJ, where images were thresholded and the percent area covered by βIII tubulin positive neurites recorded. The fold change in percent area covered was calculated for PBMC-CM treated neurons relative to control CM.

For overexpression and loss-of-function assays, *miRVana* miRNA mimics of miR-27a-3p, miR-223-3p, or Negative Control #1 (ThermoFischer); *miRVana* miRNA inhibitors miR-23a-3p, miR-27a-3p, or Negative Control #1 (ThermoFischer); and miRCURY LNA miRNA inhibitor miR-223-3p or Negative Control A (Qiagen) were used. Neurons were cultured without antibiotics and transfected at 2 DIV with a 20 nM final concentration of mimic or inhibitor using Lipofectamine RNAiMax (ThermoFischer), according to manufacturer’s instructions. miRNA inhibitors miR-23a-3p and miR-27a-3p were transfected together at a final 10 nM concentration for each inhibitor. Animal procedures for miR-223 knockout (KO) mice were approved by the Memorial University Animal Care Committee following CCAC guidelines. miR-223^−/−^ and miR-223^-/y^ KO mice (B6.Cg-*Ptprc^a^ Mir223^tm1Fcam^*/J) were purchased from Jackson Laboratories and maintained on a CD45.1^+^ Background (B6.SJL-Ptprc^a^ Pepc^b^/BoyJ) (Johnnidis *et al*., 2008). For miR-223 –/- neuronal cultures, female miR-223+/− and male miR-223-/y mice were crossed to gain WT, heterozygous and KO embryos such that each embryo was dissected individually before genotypes were confirmed. Genotyping was performed using common 5’-TTCTGCTATTCTGGCTGCAA-3’; WT 5’-CAGTGTCACGCTCCGTGTAT-3’; and KO 5’-CTTCCTCGTGCTTTACGGTATCG-3’ primers (Integrated DNA Technologies). Small molecule inhibitors for neurodegeneration assays included Cetrorelix acetate (Sigma-Aldrich), Leuprolide acetate salt (Sigma-Aldrich), GYKI 53655 hydrochoride (abcam), and MK801 hydrogen maleate (Sigma-Aldrich).

### Cell viability

Cell viability was assessed with the CellTitre-Glo (CTG) Assay (Promega) according to manufacturer’s instructions. Neurons treated with CTG Reagent were transferred to black-walled clear bottom 96-well plates, and luminescence assessed using Victor3 Plate Reader.

### Quantitative RT-PCR (qRT-PCR)

Total RNA extraction was done using miRNeasy Mini Kit, according to manufacturer’s instruction. miRNA expression was assessed using multiplex qRTPCR with Taqman miRNA Assays (ThermoFisher) and snoRNA202 as the endogenous control, as previously described (Moore *et al*., 2013; Juzwik *et al*., 2018). Fold change calculations for miRNA expression were performed using the –ΔΔCT method (Livak and Schmittgen, 2001).

### Active EAE Induction

All animal procedures related to EAE induction were approved by the Centre de Recherche du Centre Hospitalier de l’Université de Montréal Animal Care Committee following CCAC guidelines. EAE was induced in 6-9 week old female C57BL/6 mice, and clinical signs of EAE were assessed daily, as described previously (Lecuyer *et al*., 2017). Animals were immunized with subcutaneous injections of 200 μg of MOG_35–55_ (MEVGWYRSPFSRVVHLYRNGK; Alpha Diagnostic International) in 100-μL emulsion of CFA (4 mg/mL *Mycobacterium tuberculosis*; Fisher Scientific). On day 2, Pertussis toxin (500 ng PTX, Sigma-Aldrich) was injected intraperitoneally. The scoring system used was as follows: 0, normal; 1, limp tail; 2, slow righting reflexes; 2.5, difficulty walking/ataxia; 3, paralysis of one hind limb (monoparalysis); 3.5, hind limb monoparalysis and severe weakness in the other hind limb; 4, paralysis of both hind limbs (paraparalysis); 4.5, hind limb paraparalysis and forelimb weakness; 5, moribund (requires sacrifice). Animals were injected with a lethal dose of Euthanyl^®^ and perfused intracardially using cold saline for laser capture micro-dissection (LCM). For other analysis, animals were perfused with cold PBS [0.1M] followed with 4% PFA (Electron microscopy sciences).

### Laser-capture micro-dissection

The spinal cord and retina were isolated and frozen at –80°C in Tissue-Tek^®^ optimal cutting temperature (OCT) compound. Preparation of slides for LCM, LCM procedure, and extraction of RNA from laser-captured material were done as described previously(Juzwik *et al*., 2018). For LCM, lumbar motor neurons were isolated from EAE mice at naïve and peak stage; and the retinal ganglion cell (RGC) layer was isolated from EAE mice at pre-symptomatic and peak.

### Human brain samples

Snap-frozen postmortem brain samples from MS and control patients were obtained from the University Hospital Centre of Québec (Whittaker Hawkins *et al*., 2017). Biopsies were classified as acute active lesions (stage 1 and 2) or normal appearing white matter (NAWM) from secondary progressive patients and severe progressive patients. Frozen sections were cut with a cryostat to obtain approximately 20 mg of tissue, which was homogenized with QIAzol Lysis Reagent and total RNA extracted using miRNeasy Mini Kit according to manufacturer’s instructions.

### Whole mount Optic Nerve Staining and Clearing

Optic nerves with eye cups and chiasm attached were collected from EAE mice. The nerves were dissected and post-fixed for an additional 2h. We used an optimize protocol of immunolabeling-enabled three-dimensional imaging of solvent cleared organs (iDisco) to immuno-stain whole mount optic nerve. In brief, the nerves were dehydrated with increasing concentrations of methanol (MeOH, Fischer Chemical) followed by incubation with hydrogen peroxide (Fischer Chemical) overnight. The optic nerves were rehydrated again with MeOH and washed in PBS before incubation for seven days with non-phosphorylated Alexa 488 conjugated-neurofilament-H antibody (5 μg/ml per nerve; Milipore). Nerves were then washed three times with PBS, embedded in agar blocks (1%; Fisher scientific) and cleared using increasing concentration of tetrahydrofuran (THF 50%, 80% and 100%; Sigma-Aldrich) followed by immersion in benzyl ether (DBE, Sigma-Aldrich). Transparent nerves were imaged using SP8 Leica confocal microscope at the proximal, middle and distal portion of the nerve. Optic nerves were analyzed at multiple stages throughout EAE progression: presymptomatic (score 0, 8-day post-injection (dpi), onset (score 1–2.5, 12 dpi), peak (score 3 – 4.5; 14 dpi), and chronic (35 dpi).

### Retinal ganglion cell survival quantification

RGC survival was assessed on flat-mounted retina as described previously (Galindo-Romero *et al*., 2011).

### Intravitreal injection and AAV2-virus delivery

Intravitreal injections (2μl) were made in the vitreous chamber of the left eye using a custom-made glass micro-needle (Wiretrol II capillary, Drummond Scientific Co). The sclera was exposed under general anaesthesia, and the tip of the needle inserted into the superior ocular quadrant at a 45° angle through the sclera and retina into the vitreous space. AAV2-miR-223 virus (8×10^12 GC/ml) or AAV2-NT virus (AAV2-NT; 2.1×10^12 GC/ml; Penn Vector Core) were injected four weeks prior to EAE induction. qPCR analyses of miR-223 were assessed in the optic nerve and retina of injected mice for validation.

### In silico assessment of predicated targets

A bioinformatics assessment of putative target genes was performed for miR-27a-3p and miR-223-3p based on a comparative analysis by seven prediction programs for mouse transcripts. These include: Diana-microT (Reczko *et al*., 2012; Paraskevopoulou *et al*., 2013), microRNA.org (Betel *et al*., 2008), miRDP (Wong and Wang, 2015), miRTarBase (Chou *et al*., 2016), RNA22 (Miranda *et al*., 2006), TargetScan (Agarwal *et al*., 2015), and TarBase (Vlachos *et al*., 2015). Only mRNAs that were identified as putative targets across 4 of the 7 prediction programs were analyzed further, termed ‘filtered targets’(Juzwik *et al*., 2018). Pathways regulated by overlapping filtered targets were determined using an overrepresentation test by Protein ANalysis THrough Evolutionary Relationships (PANTHER) classification system (http://www.pantherdb.org/) (Mi *et al*., 2013; Mi *et al*., 2017). The p-values for the PANTHER Pathways were determined by PANTHER using the binomial statistic with a Bonferroni correction for multiple testing.

### Statistical analyses

Statistical analyses were performed using GraphPad Prism 6. As indicated in figure legends, the following statistical tests were used: student’s t-test; one-way ANOVA; two-way ANOVA; post-hocs include Dunnett’s and Tukey’s multiple comparisons tests. Sample sizes are indicated in the figure legends and significance was defined as * p<0.05, ** p<0.01, *** p<0.001, and **** p<0.0001.

## Results

### miR-27a and miR-223 are upregulated in EAE and MS

miRNA expression has been profiled in MS lesion tissue providing a rich resource of disease-relevant miRNAs (Junker *et al*., 2009). Using the EAE model, we sought to determine if candidate miRNAs from that study were regulated in inflamed neurons where they could contribute to neuronal pathology. Neurons were captured from mice subjected to EAE at peak disease by laser capture micro-dissection (LCM) and miRNA expression was analyzed by qPCR. Four miRNAs were significantly upregulated when compared to naïve control animals; miR-146a-5p (∼2 fold), miR-27a-3p (∼5 fold), miR-23a-3p (∼7 fold), and miR-223-3p (∼40 fold) (Fig. 1a). We profiled the three miRNAs with the highest fold change (miR-27a-3p, miR-23a-3p, and miR-223-3p) in the RGC layer isolated from EAE mice to see if their regulation was conserved in multiple neuronal cell types. In retinal neurons miR-27a-3p (∼7 fold) and miR-23a-3p (∼30 fold) were significantly upregulated when compared to pre-symptomatic control animals whereas expression of miR-223-3p was unaffected (Fig. 1b). To validate if these miRNAs are regulated in MS we assessed their regulation in MS tissue samples. Tissue was analyzed from post-mortem human brain samples from the normal appearing white matter (NAWM) of control and moderate MS patients, and acute active lesions of moderate and severe MS patients (Whittaker Hawkins *et al*., 2017). Acute active lesions represent early inflammatory events with initial axonal damage involving activated macrophages/microglia (Ferguson *et al*., 1997; Reynolds *et al*., 2011). By qPCR, miR-27a-3p and miR-223-3p were significantly upregulated in acute active lesions of severe MS patients compared to moderate MS and NAWM; however, in our patient cohort we did not detect the previously reported up-regulation of miR-23a-3p (Fig. 1c-e) (Junker *et al*., 2009). This may result from patient-to-patient variability. We conclude that miR-27a-3p and miR-223-3p are upregulated in neurons of EAE mice, and in tissue from active MS lesions.

**Figure 1.**
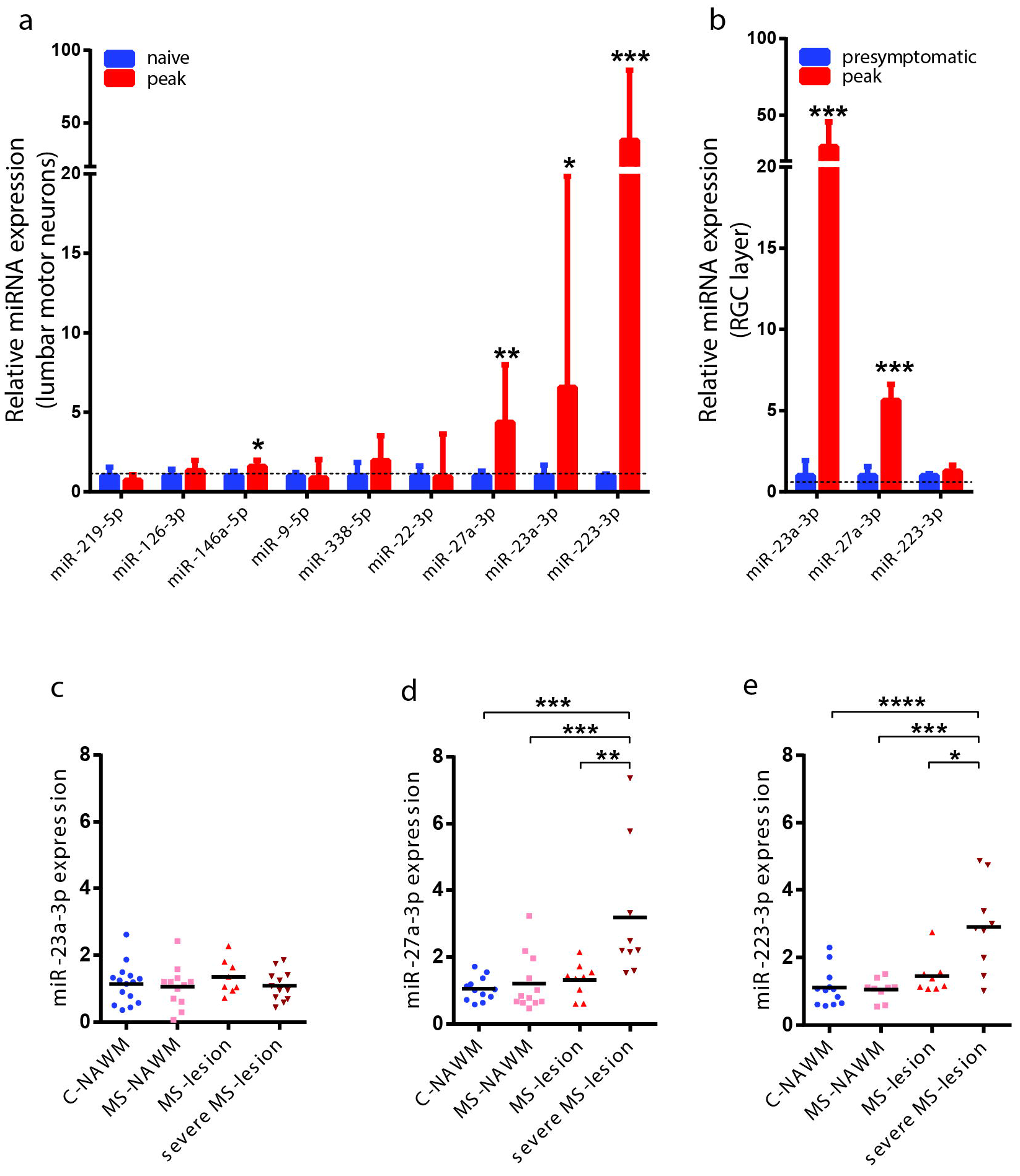
Candidate miRNA expression in EAE and MS. **(a-b)** qPCR expression data for miRNA collected from lumbar motor neurons **(a)** and retinal neurons **(b)** from EAE mice. miR-23a-3p, miR-27a-3p, and miR-223-3p are upregulated in EAE lumbar motor neurons compared to naïve mice (n = 3–6, one-way ANOVA, p<0.05, Dunnett’s post-hoc). miR-23a-3p and miR-27a-3p are upregulated in the EAE RGC layer compared to presymptomatic mice (n = 3–8, oneway ANOVA, p<0.05, Dunnett’s post-hoc). **(c-e)** qPCR expression data of miR-23a-3p **(c)**, miR-27a-3p **(d)** and miR-223-3p **(e)** in acute active lesions of severe MS patients compared to the NAWM of control and moderate MS, and the acute active lesions of moderate MS patients. (n = 7–12, one-way ANOVA, p<0.05, Dunnett’s post-hoc). qPCR data is represented as fold change..

### miR-27a-3p and miR-223-3p are neuroprotective

miR-223-3p and miR-27a-3p upregulation in EAE motor neurons correlates with motor neuron degeneration. However, previous studies have described neuroprotective roles for miR-223 and miR-27a in models of stroke and traumatic brain injury (Harraz *et al*., 2012; Cai *et al*., 2016; Sun *et al*., 2017). To test if upregulation of these miRNAs contributes to axon degeneration or compensatory neuroprotection in neurons, we performed overexpression and loss of function experiments in dissociated mouse cortical neurons. miRNA mimics for miR-27a-3p and miR-223-3p (M27a, M223) and a negative control mimic (NC) were transfected into dissociated cortical neurons at 2 DIV. Successful transfection of miRNA mimics was validated by qPCR (Supplementary Fig 2a, b). Neurons transfected with mimics for miR-223-3p and miR-27a-3p grew normally and did not exhibit any axonal swellings, breaks or other signs that one might associate with degeneration (Fig. 2a-c).

**Figure 2.**
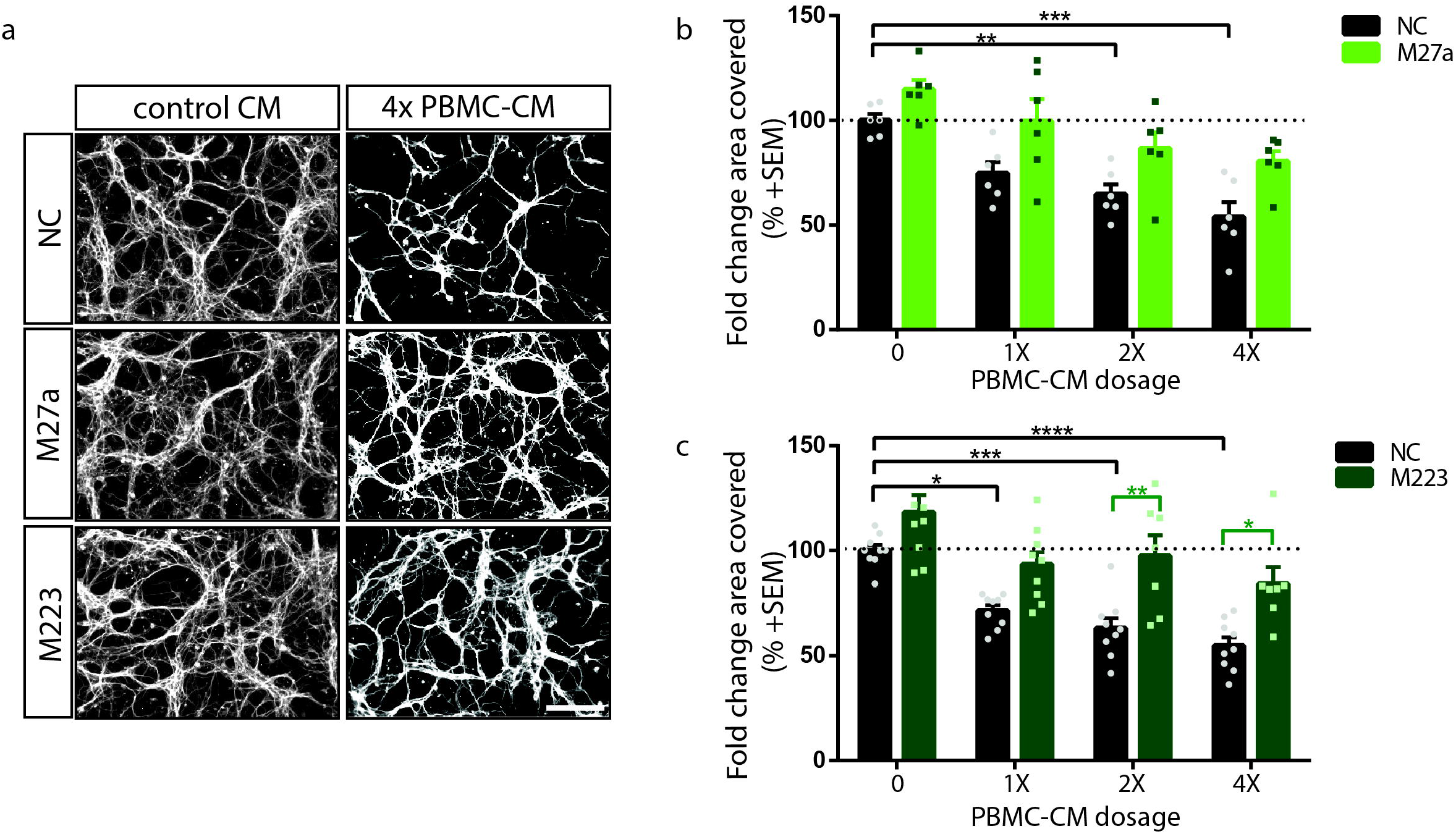
miR-27a-3p and miR-223-3p overexpression prevents PBMC-CM-dependent neurodegeneration. **(a)** βIII-tubulin-stained mouse cortical neurons transfected with miR-27a-3p, miR-223-3p, or a negative control mimic (M27a, M223, and NC, respectively) at 2 DIV and then treated for 24 h with increasing dosages of PBMC-CM at 4 DIV. **(b, c)** Quantification of the percent area covered by βIII-tubulin positive signal from thresholded images (n = 6–9, twoway ANOVA, p<0.05, Tukey’s post-hoc). Scale bar; 100 um.

We next asked if miR-223-3p and miR-27a-3p would regulate the neuronal response to MS-relevant pathological stimuli. PBMC-CM provides an excellent MS-relevant mixed stimulus containing a variety of molecules secreted from lymphocytes and monocytes. We have previously reported that CM from resting or PMA/Ionomycin (P/I)- activated PBMCs inhibits the outgrowth of dissociated neurons in culture and that this effect is mediated by secreted factors from myeloid-lineage cells (Pool *et al*., 2011; Pool *et al*., 2012). We found that dissociated cortical neurons that had been cultured at high density for 4 DIV to permit neurite outgrowth exhibited significant loss of neurite processes in the absence of cell death when treated for 12–24 hours with PBMC-CM (Fig. 2; Supp. Fig. 1). As such, PBMC-CM-dependent neurite loss represents a type of sub-lethal neurite degeneration, as opposed to a by-product of neuronal death. We then treated cortical neurons transfected with miRNA mimics with PBMCCM or control-CM. Overexpression of either miR-27a-3p or miR-223-3p protected cortical neurons from PBMC-CM-dependent neurite degeneration (Fig. 2a-c).

We hypothesized that inhibition of miR-223-3p or miR-27a-3p would sensitize neurons to the harmful effects of PBMC-CM. To test this, we transfected small, chemically modified single-stranded miRNA inhibitors that bind and inhibit endogenous miRNA molecules. Inhibition of miR-27a-3p resulted in compensatory upregulation of miR-23a, a miRNA that is a member of the same miRNA cluster, thus we co-transfected neurons with miR-27a-3p and miR-23a-3p inhibitors (Fig. 3; Supp. Fig. 2c). Inhibition of miR-27a-3p/miR-23a-3p modestly but significantly suppressed axon growth without affecting cell death consistent with the idea that this miRNA cluster may protect neurons from degeneration (Fig. 3a-c). To assess the role of miR-223, we evaluated outgrowth of dissociated cortical neurons from mir-223^−/−^ mice (Johnnidis *et al*., 2008). We also found that loss of mir-223 resulted in a significant reduction in neurite outgrowth from dissociated cortical neurons independent of cell death (Fig. 3d-f). To determine if miR-23a/27a and miR-223 loss-of-function further sensitized neurons to PBMCCM, we treated I23a/27a-transfected neurons and miR223^−/−^ neurons with PBMC-CM. I23a/27atransfected neurons were not more sensitive to PBMC-CM than NC-transfected neurons when normalized to their own basal growth (Fig. 3g). Similarly, the sensitivity of neurons from miR-223^−/−^ mice to PBMC-CM was unchanged (Fig. 3h). These results demonstrate that overexpression of miR-27a-3p and miR-223-3p is neuroprotective but that loss-of-function of the individual miRNAs is insufficient to sensitize the neurons to PBMC-dependent cytotoxicity. This may be a result of multiple miRNAs functioning in a redundant fashion to mediate neuroprotection.

**Figure 3.**
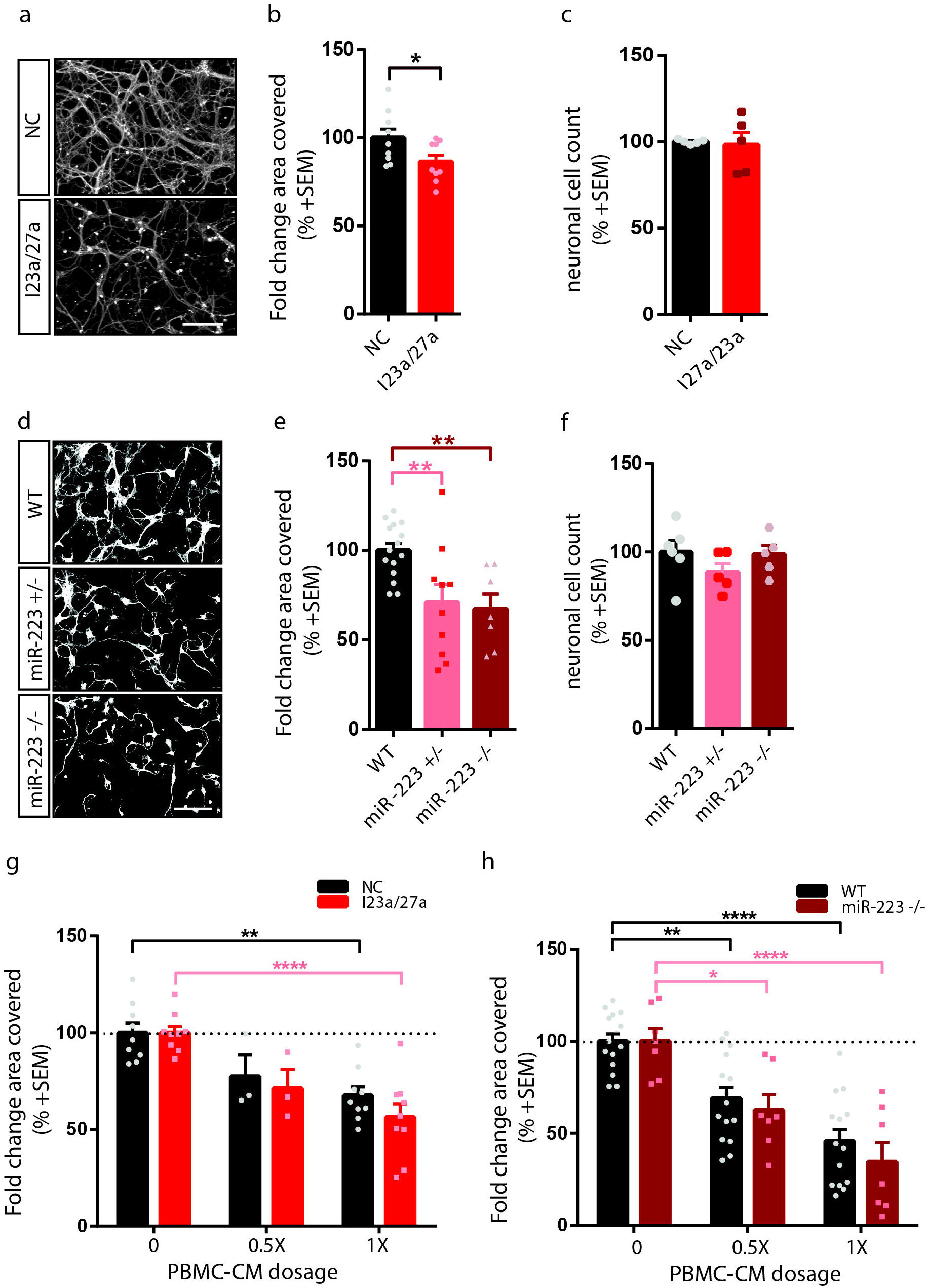
miR-27a-3p and miR-223 loss-of-function reduces basal neurite growth without further sensitization to PBMC-CM. **(a)** βIII-tubulin stained mouse cortical neurons cotransfected with miR-23a-3p and miR-27a-3p inhibitors (I23a/27a) or a negative control (NC) inhibitor at 2 DIV. **(b, c)** Quantification of neurite growth expressed as the percent of βIII-tubulin positive area in the thresholded image **(b)** or neuronal cell numbers **(c)** from I23a/27a-or NC-transfected cells **(d)** βIII-tubulin stained mouse cortical neurons from WT, miR223^+/−^ and miR223^−/−^ mice. **(e, f)** Quantification of βIII-tubulin-positive area coverage **(e)** and neuronal cell numbers **(f)** from miR223^+/−^ and miR223^−/−^ neurons. **(g, h)** Quantification of βIII-tubulin-positive area coverage from I23a/27a-and NC-transfected neurons **(g)** or miR223^−/−^ neurons **(h)** treated with PBMC-CM. Percent area covered is normalized to the respective transfection condition to determine if there is greater sensitization to PBMC-CM (n = 3–9, two-way ANOVA, p<0.05, Tukey’s post-hoc). Scale bar; 100 μm.

### miR-223 overexpression rescues axonal degeneration in EAE optic nerve

To determine if overexpression of miRNAs could also have neuroprotective activity in an *in vivo* model of CNS inflammation we chose to analyze RGCs and their axon projections in the EAE model. The optic nerve forms part of the CNS and contains the axons of RGCs, which relay visual information to the brain (London *et al*., 2013). Because of its accessibility, the visual system represents an ideal system to investigate the effect of molecules of interest in CNS. The RGCs are also readily accessible to molecular manipulation by transduction with Adeno-associated virus 2 (AAV2), which preferentially infects RGCs and produces sustained transgene expression following intravitreal injection (Cheng *et al*., 2002). Profiling of motor neurons and retinal neurons from EAE mice revealed that both miR-223-3p and miR-27a-3p were upregulated in motor neurons, whereas only miR-27a-3p was upregulated in retinal neurons (Fig. 1). We took advantage of this finding to ask if overexpression of miR-223-3p in RGCs would mediate neuroprotection *in vivo*. We optimized an iDISCO protocol to visualize axonal morphology during the progression of EAE. Optic nerves were optically cleared and immunostained with anti-neurofilament-H antibody to enable the visualization of axonal structure in 3D (Renier *et al*., 2014). Axonal swellings, ovoids and fragmentation were visible in the optic nerve throughout the progression of EAE (Fig. 4a-h). Axonal blebs were present at disease onset before significant RGC death and reached maximal density at the peak of disease (Fig. 4a-n). AAV2-miR-223 and an AAV control with a non-targeting (NT) miRNA (AAV2-NT) were injected intravitreally into the left eyes of mice four weeks prior to EAE. AAV2 was used to allow for miR-223 overexpression uniquely in neuronal cells of the retina (Davidson *et al*., 2000). Expression of miR-223 in the RGCs and optic nerve was validated by qPCR (Fig. 4o). In the contralateral optic nerve containing axons projecting from a non-injected eye, EAE induction resulted in visible focal degeneration, as evidenced by the presence of characteristic axonal blebs, ovoids, and fragmentation (Fig. 4k). In the ipsilateral optic nerve containing axons projecting from the retina injected with control AAV2-NT, a similar degree of axon degeneration was apparent (Fig. 4k). In contrast, optic nerves projecting from AAV2-miR-223-injected eyes exhibited significantly fewer axonal swellings when compared to the non-injected contralateral eye (Fig. 4l). AAV2-miR-223 injected eyes exhibited a ∼60% reduction of optic nerve degeneration (Fig. 4p). These results demonstrate that axonal swellings accumulate throughout the time course of EAE and that they can be prevented through neuron-specific miR-223 overexpression.

**Figure 4.**
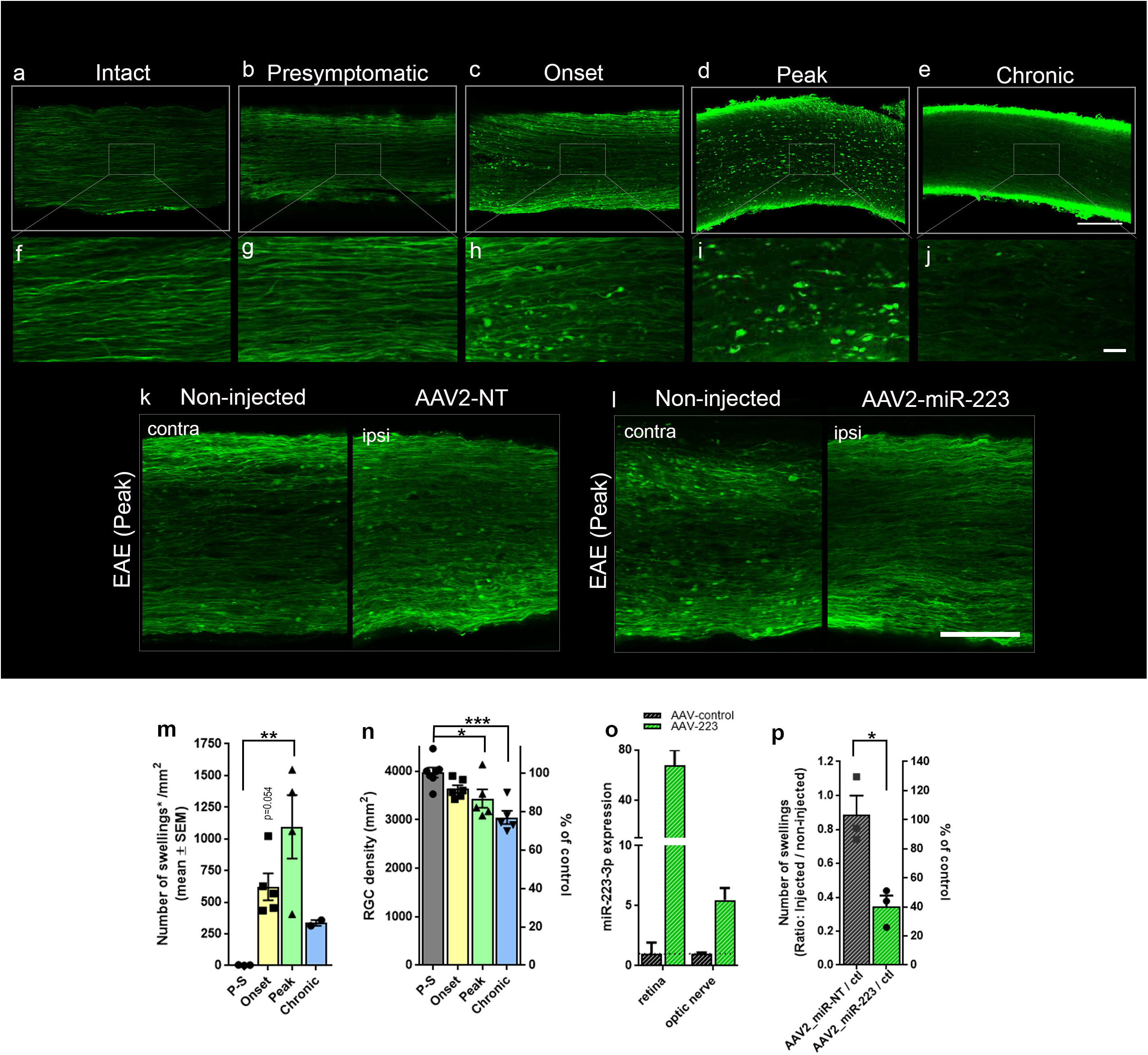
Axonal degeneration is rescued by neuronal overexpression of miR-223 *in vivo* in the EAE model. **(a-l)**, Confocal images of whole mount EAE optic nerves stained with nonphosphorylated neurofilament-H antibody showing axonal swellings occurring throughout the course of the disease in control EAE animal **(a-j)**, from AAV2-NT-injected eyes or contralateral non-injected eyes **(k)** and from optic nerves projecting from AAV-miR-223-injected eyes or contralateral non-injected eyes **(l). (f-j)** are magnifications of the boxed areas in panels **(a-e). (m)** Quantification of the density of axonal swellings in control EAE mice (n = 2–5, one-way ANOVA, p<0.05, Dunnett’s post-hoc). **(n)** Quantification of RGC survival in control EAE mice (n = 5–7, one-way ANOVA, p<0.05, Dunnett’s post-hoc. **(o)** qPCR analysis of miR-223-3p expression in retina and optic nerve collected following intravitreal injection of AAV2-miR-223 and AAV2-miR-NT (control). **(p)** Quantification of the number of axonal swellings in optic nerves following injection of AAV2-miR-223 AAV-NT expressed as a ratio of ipsi/contra; (n = 3, two-tailed student’s t-test). Ipsi = ipsilateral optic nerve that received AAV-miR treatment and contra represents the contralateral optic nerve that did not receive treatment and was thus used as an internal control. Scale bar; 100μm **(a-e)** and **(k-l)**; 15μm **(f-j)**.

### miR-27a-3p and miR-223-3p mediate neuroprotection against inflammation through the glutamate receptor (GluR) pathway

We were interested in identifying common mRNAs regulated by miR-223-3p and miR-27a-3p that are responsible for conferring neuroprotection. To identify putative mRNA targets, we used an *in silico* approach to mine predicted targets using Diana-microT(Reczko *et al*., 2012; Paraskevopoulou *et al*., 2013), microRNA.org(Betel *et al*., 2008), miRDP (Wong and Wang, 2015), miRTarBase (Chou *et al*., 2016), RNA22 (Miranda *et al*., 2006), TargetScan (Agarwal *et al*., 2015), and TarBase (Vlachos *et al*., 2015). We then retained predicted targets that occurred in at least four of seven databases. Targets were assessed for overrepresentation of pathways using PANTHER Pathway Analysis (Mi *et al*., 2013; Mi *et al*., 2017). PANTHER identified eight significantly overrepresented pathways formed by the putative targets of 27a-3p and miR-223-3p (Fig. 5a). We reasoned that miR-223-3p and miR-27a-3p may mediate neuroprotection by downregulating these signalling pathways and that inhibitors of these candidate pathways may therefore also protect neurons from PBMC-CM. We first investigated the gonadotropin hormone receptor (GnRH) pathway as it contained the most targets out of the pathways with 29 putative targets (Supplementary Table 1). GnRH receptor signaling is best described in the hypothalamic-pituitary-gonadal axis, regulating the reproductive system (Quintanar and Salinas, 2008). However, GnRH and its receptor can also be found in other cells such as the cortex and spinal cord (Albertson *et al*., 2008; Quintanar and Salinas, 2008). We treated neurons the GnRH antagonist cetrorelix or the agonist leuprolide and assessed effects on neurite outgrowth. Neither leuprolide or cetrorelix had significant effects on neurite growth (Fig. 5 b, c). Similarly, they did not significantly affect PBMC-CM induced degeneration compared to their respective controls (Figure 5b,c).

**Figure 5.**
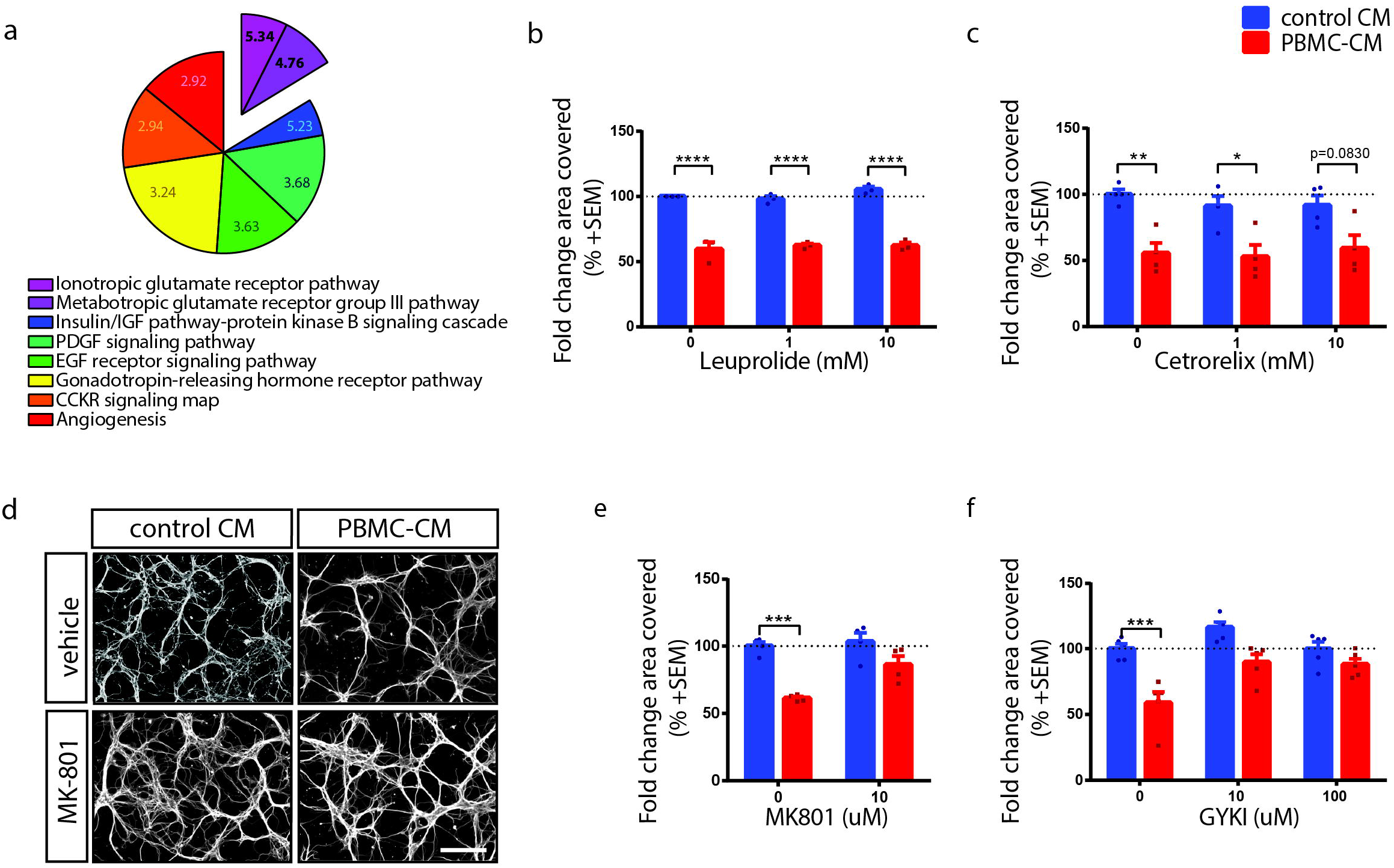
Inhibition of GluR signaling protects against neurodegeneration *in vitro*. **(a)** *In silico* identified targets of miR-27a-3p and miR-223-3p identified by a PANTHER overrepresentation test. Ionotropic GluR pathway was the most enriched pathway for miR-27a-3p and miR-223-3p targets at 5.34 fold enrichment (p = 0.004), and metabotropic glutamate receptor pathway was the third most enriched pathway at 4.76 fold enrichment (p = 0.002). All other significantly overrepresented pathways are depicted within the pie (p<0.05, Binomial Bonferroni correction for multiple testing). **(b, c)** Fold change in the percent area covered by βIII-tubulin positive neuronal cell bodies and their neurites, grown for 4 DIV and treated for 24h with PBMC-CM or control CM in the presence of GnRH agonist leuprolide **(b)** or antagonist Cetrorelix **(c)** (n = 3–6, two-way ANOVA, p<0.05, Tukey’s multiple comparisons test. ****p<0.0001, **p<0.01, *p<0.05.) **(d)** βIII-tubulin stained mouse cortical neurons in the presence or absence of MK-801 and treated with control-or PBMC-CM. **(e, f)** Quantification of fold change in percent area covered by βIII tubulin following treatment with PBMC-CM in the presence of the NMDAR antagonist MK-801 **(e)** or the AMPAR antagonist GYKI **(f)** (n = 4–5; two-way ANOVA, p<0.05, Tukey’s multiple comparisons test.

We then investigated the ionotropic glutamate receptor (GluR) pathway, as it showed the highest fold enrichment at 5.34 fold (Fig 5a). There are three main families of ionotropic glutamate receptors: AMPA receptors (AMPAR), kainate receptors, and NMDA receptors (NMDAR) (Stojanovic *et al*., 2014). To interrogate whether PBMC-CM-dependent degeneration could be blocked with ionotropic GluR antagonists, we treated cortical neurons with the NMDAR antagonist, MK-801. Notably, MK-801 prevented PBMC-CM mediated neurodegeneration relative to vehicle control (Fig. 5d,e). As NMDAR activation depends upon membrane depolarization, which is typically enacted through AMPAR activation we next sought to determine if this effect was also mediated through upstream AMPAR activation (Lau Anthony and Tymianski, 2010). Application of a small molecule inhibitor of AMPARs and kainate receptors (GYKI53655, herein GYKI) also completely inhibited neurite loss upon treatment with PBMC-CM (Fig. 5f). GYKI is predicted to inhibit greater than 90% of AMPARs at 10 uM but less than 5% of kainate receptors (Wilding, 1995). Its neuroprotective effects were therefore likely mediated through the inhibition of AMPARs as 10 uM was sufficient to prevent PBMCCM induced degeneration (Fig. 5f).

### miR-27a-3p and miR-223-3p loss-of-function inhibits neurite growth through Glu receptor regulation

We observed reduced neurite outgrowth in miR-223 and miR-27a loss-of-function experiments and thus we hypothesized that this might be a result of increased GluRs on the neuronal cell surface that would sensitive neurons to low concentration of glutamate in the media. Consistent with this model we found that MK-801 and GYKI were both able to rescue LNA-mediated miR-223-3p loss-of-function, restoring growth to that of control neurons (Fig. 6 a-c). Similarly, MK-801 and GYKI were also able to rescue miR-23a-3p/27a-3p loss-of-function (Fig. 6 d-f). Overall, our results suggest that overexpression of miR-223-3p and miR-27a-3p downregulate ionotropic GluRs to promote neuroprotection.

**Figure 6.**
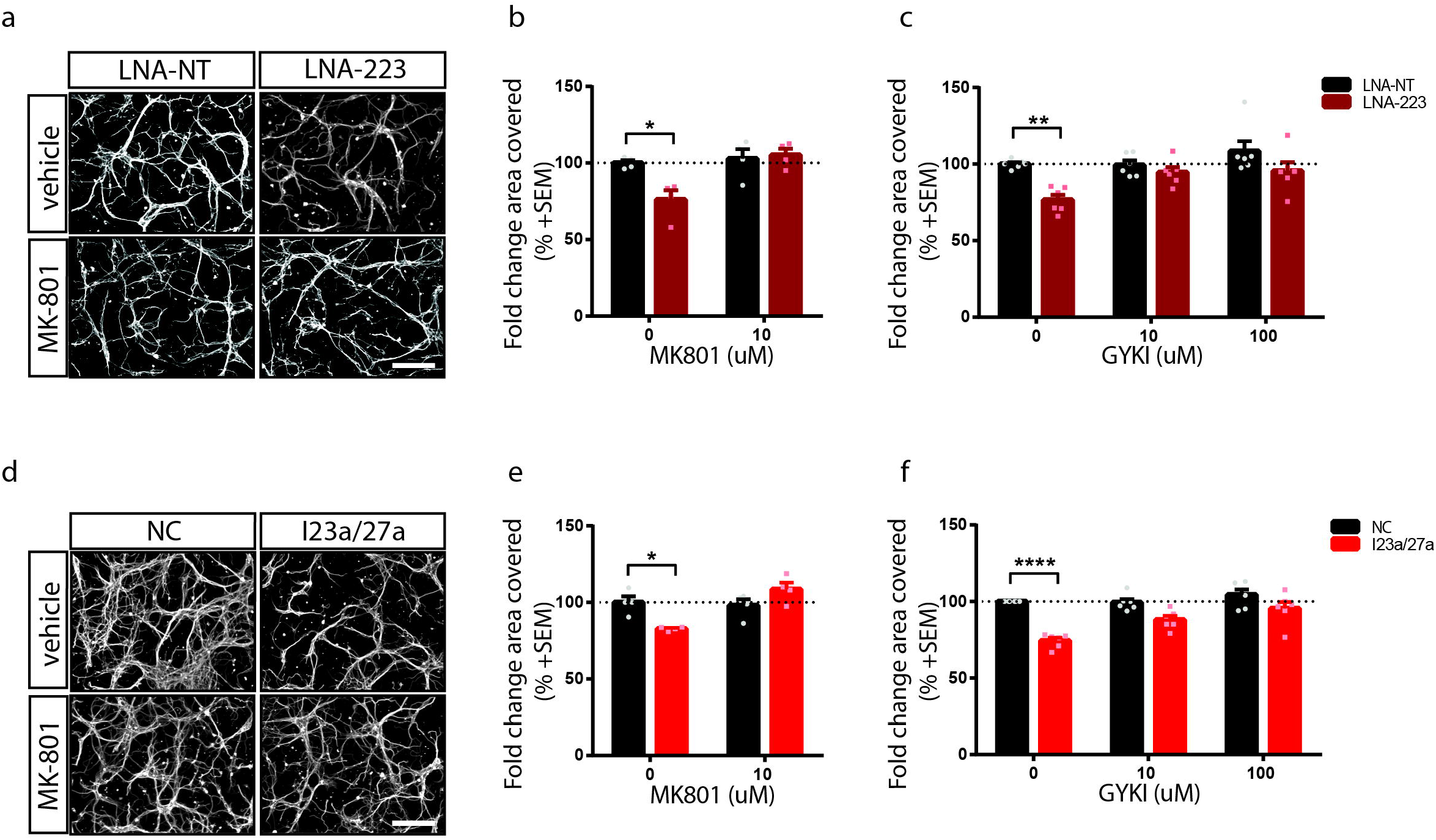
miR-27a-3p and miR-223-3p loss-of-function growth deficits can be restored through GluR inhibition. **(a,d)** βIII-tubulin-positive cortical neurons transfected with miR-223-3p LNA inhibitor or LNA non-targeting control (**a**, LNA-223, LNA-NT) or miR-23a-3p and miR-27a-3p inhibitors or negative control inhibitor (**d**, I23a/27a, NC) in the presence of MK-801. **(b, c, e, f)** Quantification of the fold change area covered by βIII tubulin in miR-223 **(b, c)** and I23a/27a **(e, f)** knockdown conditions in the presence of MK801 **(b, e)** or GYKI **(c, f).** Neurons were transfected with miRNA inhibitors at 2 DIV and treated at 4 DIV for 24h with MK-801 or GYKI (n = 4–6, two-way ANOVA, p<0.05, Tukey’s multiple comparisons test). Scale bar; 100μm.

## Discussion

There is good evidence that neuronal pathology underlies progressive forms of MS and neurological disability in the disease (Chiaravalloti and DeLuca, 2008). MRI and histological studies demonstrate the occurrence of axonal transection early in the disease process and have revealed that the level of axonal loss positively correlate with poor clinical scores (De Stefano *et al*., 1998; Trapp *et al*., 1998a; Wujek *et al*., 2002). Brain atrophy, reflective of cell death, is also a significant feature of progressive MS (Bermel and Bakshi, 2006). Our results describe this characteristic neurodegeneration in the optic nerve in EAE, which can be ameliorated through overexpression of miR-223. We propose this effect is mediated in part through the suppression of glutamate excitotoxicity.

A multitude of signals including cytokines, complement, free radicals, nitric oxide, proteases, excess glutamate and calcium contribute to a deleterious environment for neurons in MS (Trapp *et al*., 1998b; Yong *et al*., 2007; Gonsette, 2008). However, recapitulation of this environment *in vitro* has proved challenging. To capture this complex environment *in vitro*, we used PBMC-CM treatment. Neurons grown in the presence of PBMC-CM undergo loss of neurites, a process independent of cell death (Pool *et al*., 2011; Pool *et al*., 2012). PBMC-CM has been shown to contain glutamate in the 100 uM range (O’Driscoll *et al*., 2013). Glutamate is an excitatory neurotransmitter and amino acid, which also plays a role in metabolism (Stojanovic *et al*., 2014). Elevated glutamate levels have been shown in various tissues of MS patients (Levite, 2017). High levels of glutamate can potentiate excitotoxicity, a neurotoxic mechanism primarily activated by the influx of excess Ca^2+^ through NMDARs leading to activation of intracellular Ca^2+^-dependent caspases, breakdown of cytoskeletal components, accumulation of reactive oxygen species, and energetic failure leading to death in mature neurons (Lau Anthony and Tymianski, 2010). Immature neurons however are resistant to glutamate toxicity. Rather, treatment of immature neurons with excess glutamate results in a significant loss of dendritic arbors *in vitro* without cell death or a decrease in viability (Monnerie *et al*., 2003). In a similar manner, miR-27a-3p and miR-223-3p overexpression also rescued the phenotype observed in PBMC-CM treated cultures. Conversely, their inhibition resulted in basal growth deficits. These basal growth deficits could be rescued by application of either MK801 or GYKI. Based on this evidence, we propose both miRNAs to be involved in regulating components of the ionotropic GluR pathway. This is consistent with previous findings demonstrating that miR-223 directly targets GluR2 and NR2B and mediates neurotprotection in a stroke model(Harraz *et al*., 2012) Additionally, we suggest that inhibition of either miRNA sensitizes neurons to extracellular glutamate, through their proposed afore-mentioned role. PBMC-CM treatment in the presence of the inhibitors did not exacerbate the phenotype, suggesting that GluRs may have been saturated prior to the addition of PBMC-CM.

Both the miR-23a∼27a∼24-2 cluster (which produces miR-27a) and miR-223 represent miRNAs with previously characterized roles in regulating the immune system in MS and EAE. miR-27a drives a pro-inflammatory phenotype in macrophages and promotes the expression of pathogenic Th17 cells in MS (Xie *et al*., 2014; Ahmadian-Elmi *et al*., 2016). Similarly, miR-223 promotes pro-inflammatory macrophages and Th1/Th17 differentiation. This is emphasized in miR-223 KO mice, which have reduced EAE disease severity linked to decreased inflammation (Ifergan *et al*., 2016; Satoorian *et al*., 2016; Cantoni *et al*., 2017). However, very little was previously known about their roles exclusively in the neuronal compartment. The retina and optic nerve in EAE represented an attractive model of CNS degeneration to study due to its accessibility and relevance to MS (Green *et al*., 2010; London *et al*., 2013). We described the accumulation of axonal swellings in the optic nerve over the time course of EAE using an optimized protocol combining 3DISCO and iDISCO. Using AAV2-mediated overexpression of miR-223, we successfully manipulated miR-223 expression *in vivo*, allowing us to study its role specifically within neurons in disease. We chose to overexpress miR-223 as opposed to miR-27a in the RGCs as miR-223 was not intrinsically upregulated in RGCs in EAE. Here we saw a robust neuroprotective effect of miR-223 where its overexpression led to a considerable reduction of axonal damage in the optic nerve in EAE. This suggests that miR-223 has an important and protective role within neurons in addition to its well described immunomodulatory roles.

Our *in silico* analysis identified *Gria2* and *Grin2b* as putative targets of miR-27a-3p and miR-223-3p (Supplementary Table 1). Harraz and colleagues previously validated *Gria2* and *Grin2b* as bona fide targets of miR-223-3p in rat hippocampal neurons through luciferase assay and western blot(Harraz *et al*., 2012). Overexpression of miR-223 also directly inhibited NMDA-induced Ca^2+^ influx *in vitro* and *in vivo* enhancing neuroprotection in a model of stroke (Harraz *et al*., 2012). We suspect that AAV2-mediated overexpression of miR-223 may be likewise potentiating neuroprotection through the inhibition of NMDAR mediated Ca^2+^ influx, mitigating glutamate excitotoxicity mediated neurodegeneration during EAE. *Gria2*, an AMPAR subunit, is another validated target of miR-223-3p. Downregulation of *Gria2* has been shown to lead to less efficient AMPAR assembly, contributing to an overall reduction of AMPARs at the synapse (Sans *et al*., 2003). *Gria2* targeting by miR-223-3p may therefore contribute to a decrease in AMPAR expression. As glutamate is produced and released in large quantities by activated immune cells which are known to congregate at active demyelinating lesions in MS (Kostic *et al*., 2013), a global decrease in AMPARs and a concomitant reduction of NMDAR mediated Ca2+ influx by miR-223-3p-mediated gene silencing may effectively diminish the toxicity of increased glutamate present in the extracellular milieu in EAE.

In sum, we describe upregulation of miR-27a-3p and miR-223-3p in human MS lesions and characterized their expression in neurons from EAE. Previous studies have used GluR antagonists to treat EAE: treatment with either the AMPAR/kainate receptor antagonist NBQX or NMDAR antagonist, memantine, resulted in a decreased clinical score and reduced axonal damage (Wallstrom *et al*., 1996; Pitt, 2000; Kostic *et al*., 2013). Likewise, we demonstrated that overexpression of miR-223 is neuroprotective in EAE which we propose to be through its similar ability to target components of the ionotropic GluR pathway reducing glutamate excitotoxicity in the disease. Although validation of the miR-27a-3p targets within the GluR pathway has not been conducted, based on our *in vitro* experiments we propose that both miRNA converge on this pathway to exert a neuroprotective effect. Future experiments will be able to show whether miR-27a-3p has direct functional targets within the GluR pathway, and whether its overexpression *in vivo* may also mitigate axonopathy in EAE. Nevertheless, our results here demonstrate a strong case for miRNA-mediated silencing of GluRs as a powerful tool in preventing axonal degeneration in EAE.

## Acknowledgements

The authors would like to acknowledge Drs. Nicolas Paradis-Isler and Matthew Parsons for helpful discussions on the manuscript and Valina Dawson for providing the AAV2-miR-223 plasmid. AEF is funded by the Canadian Insitutes for Health Research and the Multiple Sclerosis Society of Canada (MSSOC). BM, CAJ, SSD and YZ were funded by awards from the MSSOC/Fonds de recherché du Quebec, Vanier Canada Graduate Scholarships, Canada Graduate Scholarship – master’s award allocation and MSSOC, respectively.

## Abbreviations

3’ untranslated region: (3’UTR)
AMPA receptor: (AMPAR)
benzyl ether: (DBE)
Canadian Council on Animal Care: (CCAC)
CellTitre-Glo: (CTG)
central nervous system: (CNS)
days in vitro: (DIV)
days post injection: (dpi)
embryonic day: (E)
experimental autoimmune encephalomyelitis: (EAE)
fetal bovine serum: (FBS)
glutamate receptor: (GluR)
gonadotropin-releasing hormone: (GnRH)
immunolabeling-enabled three dimensional imaging of solvent cleared organs: (iDISCO)
knockout: (KO)
laser capture microdissection: (LCM)
messenger RNA: (mRNA)
methanl: (MeOH)
microRNA: (miRNA)
multiple sclerosis: (MS)
Multiwavelength Cell Scoring: (MWSC)
NMDA receptor: (NMDAR)
optimal cutting temperature: (OCT)
optic nerve: (ON)
paraformaldehyde: (PFA)
peripheral blood mononuclear cell: (PBMC)
PBMC conditioned media: (PBMC-CM)
poly-l-lysine: (PLL)
primary progressive multiple sclerosis,: (PPMS) polymerase chain reaction (PCR)
primary progressive multiple sclerosis: (PPMS)
Protein ANalysis THrough Evolutionary Relationships: (PANTHER)
quantitative RT-PCR: (RTqPCR)
relapsing-remitting multiple sclerosis: (RRMS)
retinal ganglion cell: (RGC)
tetrahydrofuran: (THF)
wild type: (WT)

